# A live cell protein complementation assay for ORFeome-wide probing of human HOX interactomes

**DOI:** 10.1101/2022.11.09.515750

**Authors:** Yunlong Jia, Jonathan Reboulet, Benjamin Gillet, Sandrine Hughes, Christelle Forcet, Violaine Tribollet, Nawal Hadj Sleiman, Cindy Kundlacz, Jean-Marc Vanacker, Françoise Bleicher, Samir Merabet

**Affiliations:** IGFL, CNRS UMR5242, ENS-Lyon, UCBL-1, INRA USC1370, 32 Av. Tony Garnier, 69007 Lyon, France; Department of Developmental and Cell Biology, University of California, Irvine, Irvine, CA 92697, USA; LiPiCs, 46 Allée d’Italie, 69007 Lyon, France

## Abstract

Biological pathways rely on the formation of intricate protein interaction networks called interactomes. Getting a comprehensive map of interactomes implies developing tools that allow capturing transient and low affinity protein-protein interactions (PPIs) in live conditions. Here we present an experimental strategy, Cell-PCA (Cell Protein Complementation Assay), which is based on BiFC (Bimolecular Fluorescence Complementation) and high throughput sequencing for ORFeome-wide analyses of different interactomes in the same live cell context. The specificity and sensitivity of Cell-PCA was established by using a wild type and a single amino-acid mutated HOXA9 protein, and the approach was subsequently applied for seven additional human HOX proteins. These proof-of-concept experiments revealed novel molecular properties of HOX interactomes and led to the identification of a novel cofactor of HOXB13 for promoting its proliferative activity in a cancer cell context. Taken together, our work demonstrates that Cell-PCA is pertinent for revealing and, importantly, comparing interactomes between different or highly related bait proteins in the same cell context.

## Introduction

Organismal development and fitness depend for a large part on their cell protein content. Proteins are versatile molecules that work in a crowded environment, establishing a number of interactions with other surrounding proteins. These protein-protein interactions (PPIs) form intricate networks, called interactomes, which are changing from cell-to-cell and stage-to-stage.

Characterizing these dynamic molecular networks is a key issue for understanding protein function and implies developing highly sensitive tools. One main experimental approach to capture PPIs is the yeast two-hybrid system (Y2H), which relies on the indirect readout of a reporter gene to reveal any interaction between a bait protein and candidate partners (Paiano *et al*., 2019). Although very popular, Y2H presents the major inconvenient of being performed in a heterogeneous context (i.e., in the yeast for non-yeast proteins and in the context of proteins fused to heterologous DNA-binding and activation domains).

Recent approaches based on biotin-ligase enzymes and Liquid Chromatography-Mass Spectrometry (LC-MS) identification of biotinylated partners constitute promising alternatives for the characterization of interactomes in specific cell- and tissue types (Varnaitė & MacNeill, 2016; Carnesecchi *et al*, 2020). These approaches are appropriate for the identification of endogenous interactomes but are not compatible for more systematic and high throughput interrogations of binary PPIs with dedicated candidate libraries.

To date, the use of fluorescent (Miller *et al*, 2015) and non-fluorescent (*Remy et al*., 2007) reporters for protein-fragment complementation assays (PCAs) represent the simplest and most sensitive tools for screening PPIs in live cell conditions. These approaches rely on the property of the reporters to be reconstituted from separate N- and C-terminal fragments upon spatial proximity (Fig. 1A). However, all currently described fluorescent PCA-based screens relied on Open Reading Frames’ libraries (ORFeomes) that were transiently used in the cell, either upon co-transfection (Co-PCA strategy: Fig. S1 and Remy & Michnick, 2004; Berendzen et al., 2012) or transduction of lentiviral particles (Re-PCA strategy: Fig. S1 and Ding *et al*, 2006; Lee *et al*, 2011; Simon E. Coopera, 2015). These strategies do not allow comparing interactomes between different bait proteins since the libraries were not stably conserved and were systematically screened in one shot with no replicates in the same cell population.

**Figure 1.**
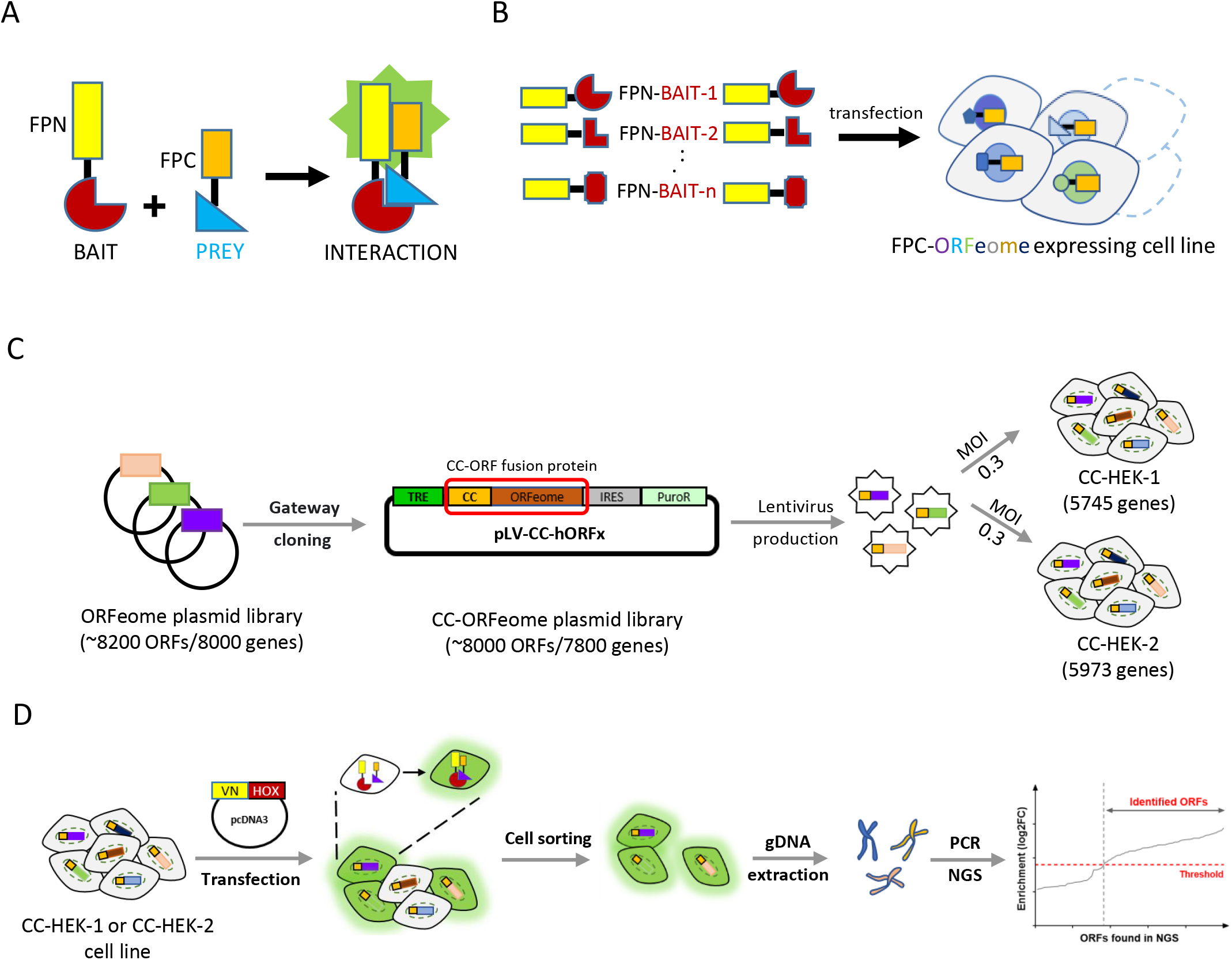
Principle of the Protein Complementation Assay (PCA) and its applications in large-scale interaction screens. **A**. Application of fluorescent-based PCA for revealing interaction between two candidate partners. The N (FPN)-or C (FPC)-terminal fragment of the fluorescent protein (FP) is fused to one of the two putative interaction partners (bait and prey proteins). The interaction between the bait and the prey protein allows the reconstitution of the fluorescent protein and the emission of fluorescent signals upon excitation. This principle of complementation has also been developed with enzymes for large-scale interaction screens (see for example (*Remy et al*., 2007)). **B-** Application of PCA-based strategies for large-scale interaction screens in living cells. The cell line PCA-based screening strategy (Cell-PCA) relies on the use of cell lines established with an inserted FPC-fusion library (D). These cell lines can be used multiple times for screening for interacting partners of different FPN-fusion bait proteins introduced by transfection. **C-D**. Experimental procedure for the Cell-PCA-based screen. **C**. A pool of ~8200 human ORFs derived from the hORFeome *v3*.*1* was cloned *en masse* by Gateway® LR reaction into the lentiviral vector pLV-CC (Fig. S1), subsequently generating the CC-ORFeome plasmid library (pLV-CC-hORFs). The final expression plasmid library (~8000 ORFs) was used to produce lentiviruses and infect two different batches of HEK293T cells to generate two different cell lines (CC-HEK-1 and CC-HEK-2). **D**. Each CC-HEK cell line can be transfected with the VN-HOX-encoding plasmid. Any interaction with a CC-ORF leads to fluorescent cells that are collected using flow cytometry. Genomic DNA (gDNA) is extracted from the fluorescent sorted cells and interacting ORFs are identified through a next generation sequencing (NGS) dedicated approach. CC: C-terminal fragment of mCerulean (residues 155-238). VN: N-terminal fragment of mVenus (residues 1-172). MOI: multiplicity of infection.

Here, we propose an alternative cell line PCA-based experimental strategy (Cell-PCA) that relies on the establishment of a cell line expressing a PCA-compatible human ORFeome (Fig. 1B). This cell line can be amplified and used several times to simultaneously screen for interactions of different bait proteins, eventually allowing comparing interactome properties against the same ORFeome in the same cell context (Fig. 1B).

As a proof-of-concept, we applied this novel experimental strategy to the HOX protein family, which is involved in the regulation of numerous processes during embryonic development (Pearson *et al*, 2005; Moisés Mallo, 2018) and adult life (Seifert *et al*, 2015). HOX proteins are transcription factors (TFs) and therefore act by regulating the expression of downstream target genes *in vivo*. Several cofactors have been identified for different individual HOX proteins in various cell and developmental contexts (Carnesecchi *et al*, 2020; Lambert *et al*, 2012; Baeëza *et al*, 2015; Bischof *et al*, 2018), but no systematic large-scale interaction screening has been performed for several HOX members in the same biological system. As a consequence, very little is known about their general and specific interactome properties. For example, the question of HOX cofactor specificity remains poorly understood: is there a large proportion of specific versus common cofactors between different HOX proteins? Along the same line, work with mouse HOXA1 showed that a number of cofactors were not traditional TFs, suggesting that HOX proteins could act at different regulatory levels (Lambert *et al*, 2012). Whether this property could apply more largely to other HOX protein members remains to be investigated.

To tackle the issue of HOX interactome properties, we present the first large-scale screening of PPIs for eight different human HOX proteins in the same cell population. Our results showed that TFs are generally not employed as HOX-specific cofactors, but instead used in specific combinations in the different HOX interactomes underlying common biological functions. In contrast, we observed that HOX proteins had a general propensity to interact with non-TFs and that these interactions are more HOX-specific than the interactions with TFs. Several of these interactions were also individually validated by BiFC and co-immunoprecipitation experiments. Finally, we revealed a novel interaction that is important for HOXB13 proliferative activities in a prostate cancer-derived cell line.

Taken together, our results establish Cell-PCA as an innovative and promising experimental strategy for assessing the issue of interactome specificity in a live cell context.

## Results

### Cell-PCA screen design

The fluorescence-based complementation approach, also called BiFC (Bimolecular Fluorescence Complementation), relies on the property of hemi-fragments of monomeric fluorescent proteins such as the GFP (Green Fluorescent Protein) or Venus, to reconstitute a functional fluorescent protein upon spatial proximity (Miller *et al*, 2015). For the Cell-PCA design, we used a library of 8200 ORFs and fused this set of ORFs to the C-terminal fragment of the blue fluorescent protein Cerulean at the 5’ end (fragment CC, Fig. 1C and Materials and Methods). This fragment can complement with the N-terminal fragment of Venus (VN, leading to a Venus-like fluorescent signal) or Cerulean (CN, leading to a Cerulean-like fluorescent signal), enabling to simultaneously visualize interactions of two different bait proteins with a common cofactor (Dard *et al*, 2018a; Bischof *et al*, 2018; Hu & Kerppola, 2003).

The CC-ORF library was cloned in a lentivirus vector, downstream of regulatory sequences (*Tet-Responsive Element* (*TRE)*, Fig. 1C and Fig. S2) that respond to the tTA (tetracycline-controlled transactivator) factor in the presence of Doxycycline. The CC-ORF pooled plasmid library was used for producing lentiviral particles and subsequent infection of HEK-293T cells (see Materials and Methods). Referring to the functional titer, the pooled lentiviral libraries were transduced at a low multiplicity of infection (MOI) to achieve only one stably-integrated CC-ORFeome in most cells (see Materials and Methods). The operation was repeated two times and the two resulting cell lines were named CC-HEK-1 and CC-HEK-2, which respectively encompasses 5745 and 5973 genes (Fig. 1C and Dataset EV1-2). The basal expression of the CC-ORFeome library was further verified in the established CC-HEK cell lines by immunostaining against the CC fragment (Fig. S2).

Each VN-bait fusion protein was under the control of the constitutive *CMV* promoter (see Materials and Methods). Transfecting the VN-fusion plasmid into the established CC-HEK cell line will lead to BiFC-positive signals only when interaction occurs between the VN-bait and the CC-prey protein produced from the corresponding integrated CC-ORF (Fig. 1D). The fluorescent cells are then sorted using flow cytometry, from which the genomic DNA is extracted to prepare a sequencing library with specific oligonucleotides matching the CC-ORF construct (Fig. 1D, Fig. S3 and Materials and Methods). The presence and relative abundance of the integrated CC-ORFs in sorted fluorescent cells were assessed using a dedicated next-generation sequencing (NGS) approach (see Materials and Methods). This targeting approach allows to reduce the sequencing effort to only the beginning of the inserted CC-ORF, instead of the complete genome or insert fragments (that are highly variable in size). This strategy improves the sequencing coverage together with a reduced cost.

### Proof of concept Cell-PCA screen for HOXA9 interactomes

As a proof-of-concept, we performed a high throughput interaction screen with the human HOXA9 protein, whose interaction with two known cofactors, PBX1 and MEIS1, has been extensively described by using BiFC in HEK293T and other cell lines (Dard *et al*, 2019a). This previous work established BiFC as a specific and sensitive method for deciphering HOXA9 interaction properties, therefore establishing the appropriateness of our tools in the context of a large-scale BiFC interaction screen. We also considered a mutant form of HOXA9 as a supplementary control of the experimental design (VN-HOXA9^w^). This form is mutated in a unique conserved Trp residue that mediates the interaction with the PBX cofactor in the context of HOX/PBX dimeric complexes (Dard *et al*, 2019b; LaRonde-LeBlanc and Wolberger, 2003). This conserved Trp residue was also shown to have additional and versatile activities that changed depending on the cell context, suggesting that it could interact with other cofactors (Dard *et al*, 2019b). This Trp-mutated form of HOXA9 was therefore considered as a good control for assessing the specificity and sensitivity of our experimental tools.

The pilot screens with VN-HOXA9 and VN-HOXA9^W^ were sequentially performed in a single replicate in each CC-HEK-1 and CC-HEK-2 cell line for two main reasons. First, as previously mentioned, transduction was performed with a low MOI to get only one CC-ORF construct in the majority of cells (see Materials and Methods). This transduction condition resulted in incomplete integration of the CC-ORFeome library in each cell line (around 70%). Performing the screen in the two CC-HEK cell lines increases the proportion of the human ORFeome that could be considered (reaching 82%). Second, given the number of the VN-HOX fusion proteins used subsequently in our screen, we wanted to assess whether the strategy of considering interactions that were either specific of one CC-HEK cell line or common to the two CC-HEK cell lines could be pertinent. The two CC-HEK cell lines were thus used as biological replicates for assessing the reproducibility of interactions when considering their large pool of commonly integrated CC-fusion genes (Fig. 2A). Interactions that were specific to a cell line or common to the two cell lines were selected by applying different threshold criteria (see Materials and Methods).

**Figure 2.**
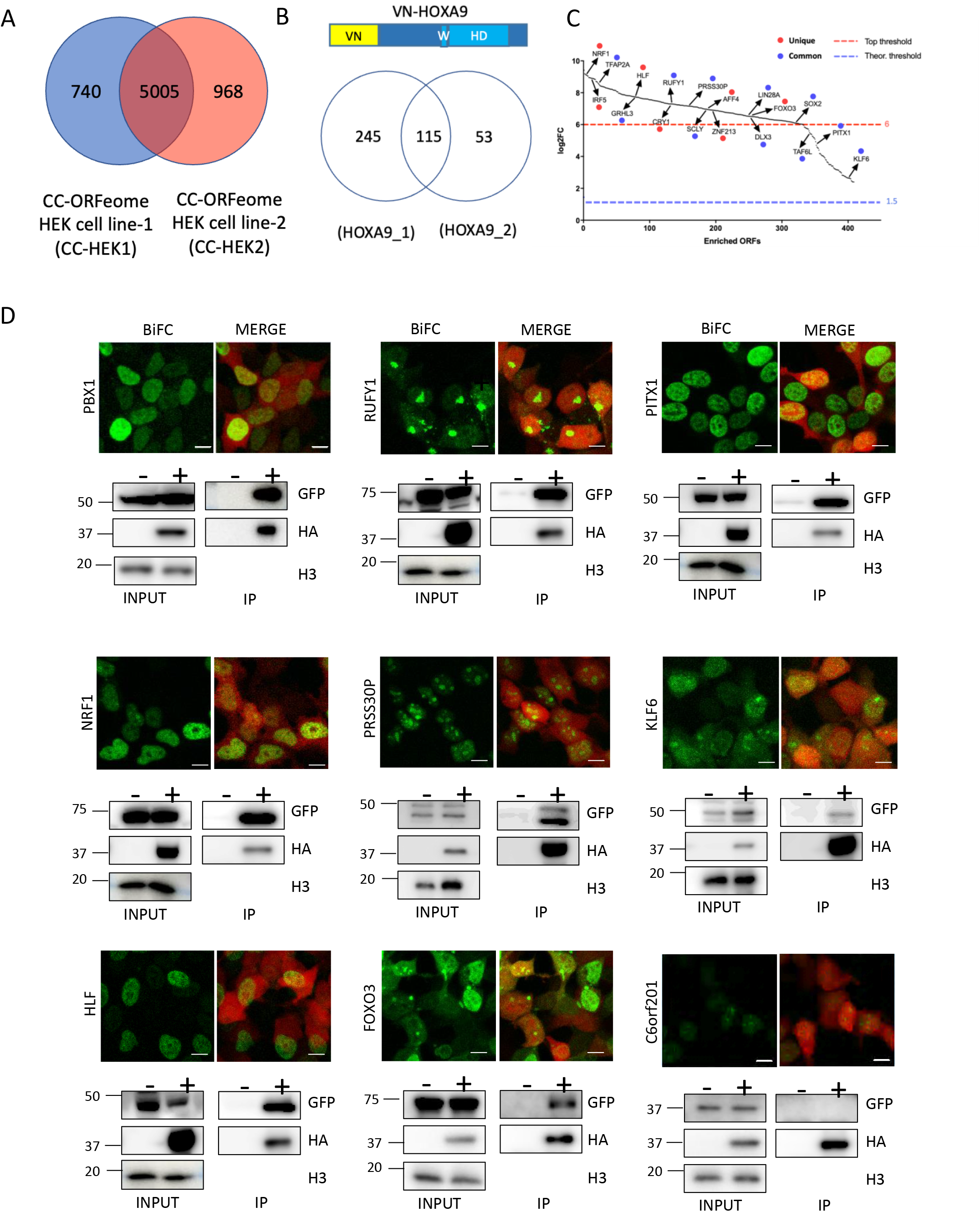
Establishing the Cell-PCA screening strategy with HOXA9. **A**. Venn diagram depicting the number of integrated ORFs in the CC-HEK-1 (blue) and CC-HEK-2 (red) cell lines. **B**. Venn diagram showing the number of HOXA9-positive ORFs in the two CC-HEK cell lines. VN-HOXA9 is schematized above the Venn diagram (with the Trp-containing motif -W- and the homeodomain -HD-). **C**. Plot of the 413 selected HOXA9-interacting candidates, ranked from the most to the lowest enriched in the Cell-PCA assay. Among them, 18 CC-ORFs were randomly picked for individual validation by BiFC, using two criteria: the 7 red dots were unique to one CC-HEK cell line with a log2 fold change (FC) superior to 6; the 11 blue dots were common and revealed in the two CC-HEK cell lines with a log2FC superior to the background (1,5). See also Materials and Methods. **D**. Illustrative confocal pictures of BiFC and co-immunoprecipitation (Co-IP) in HEK293T cells between HOXA9 and the selected candidates, as indicated. All pictures are illustrative of two independent biological replicates. For BiFC experiments, the mCherry reporter (red, merge panels) is indicative of the transfection efficiency. Note the various intra-cellular BiFC profiles with the different candidates. BiFC with the negative C6orf201 control was also negative in BiFC (with a fluorescent signal below 15% of the fluorescence intensity resulting from HOXA9/PBX1 BiFC on average) and co-IP experiments. Scale bar, 10μm. Co-IP was performed with HA-HOXA9 and the CC-ORF was revealed with anti-GFP. Staining with Histone H3 antibody validates the correct protein extraction in each condition. “+” and “−” respectively denote the co-transfection or not of HA-HOXA9 with the CC-ORF construct. The protein size scale is indicated on the left side (KDa).

The basal expression of the TRE promoter was used for the BiFC screen since it was sufficient to reveal the expression of the integrated CC-ORFs (Fig. S2). In addition to simplify the protocol, having a minimum expression level for each integrated CC-ORF also allows performing the screen in more stringent conditions without overexpression of the corresponding prey protein. Under these conditions, fluorescent signals were only observed upon co-transfection of VN-HOX9 (Fig. S4), and this pattern was systematically obtained in the subsequent screens.

According to our experimental protocol and selection criteria, we found a total of 413 (6% of the integrated CC-fusion ORFeome) positive interactions, among which 115 were common to the two CC-HEK cell lines (Fig. 2B and Dataset EV3). These candidates were selected after applying a log2 fold change enrichment threshold that was different depending on the positive interaction status in one or the two CC-HEK cells (Fig. 2C and Materials and Methods). To assess whether our post-NGS selection criteria were correct for unique or duplicated positive interactions, we randomly selected 18 CC-ORFs with a broad range of enrichment scores and that were positive either in one (7 CC-ORFs) or the two CC-HEK cell lines (11 CC-ORFs). The interaction with PBX1, which was not present in our CC-ORF libraries, was used as a positive control. We also considered one CC-ORF (C6orf201) that was negative in the screen. These candidates were tested by doing individual BiFC with VN-HOXA9, using the interaction with PBX1 as a calibrator of microscope parameters for repeating and comparing individual BiFC assays between the different biological replicates (see Materials and Methods). Individual interactions were also tested by performing co-immunoprecipitation experiments with HA-tagged HOXA9, therefore independently of the complementation system. Results showed that the interaction status was confirmed for all tested candidates (Fig. 2D and S5). Interestingly, BiFC analyses revealed various interaction profiles in live cells (Fig. 2D and S5). Collectively, these observations confirmed that the applied filtering criteria were appropriate for selecting positive interactions when considering either one or the two cell lines.

We next performed the ORFeome-wide BiFC screen in the two CC-HEK cell lines with VN-HOXA9^W^. This mutated form of HOXA9 had fewer positive interactions than wild type HOXA9 (342 in total, corresponding to 5% of the integrated CC-fusion ORFeome, and using identical selection criteria as for HOXA9: Fig. 3A and Dataset EV4). 96/413 (23%) of HOXA9 interactions were also found with HOXA9^W^, indicating that our tools were sensitive enough to reveal different interactomes upon a single amino acid mutation (Fig. 3B). The observation that 246 interactions were specific of HOXA9^w^ also highlight that the Trp mutation induced more a gain than a loss of the HOXA9 interaction potential.

**Figure 3.**
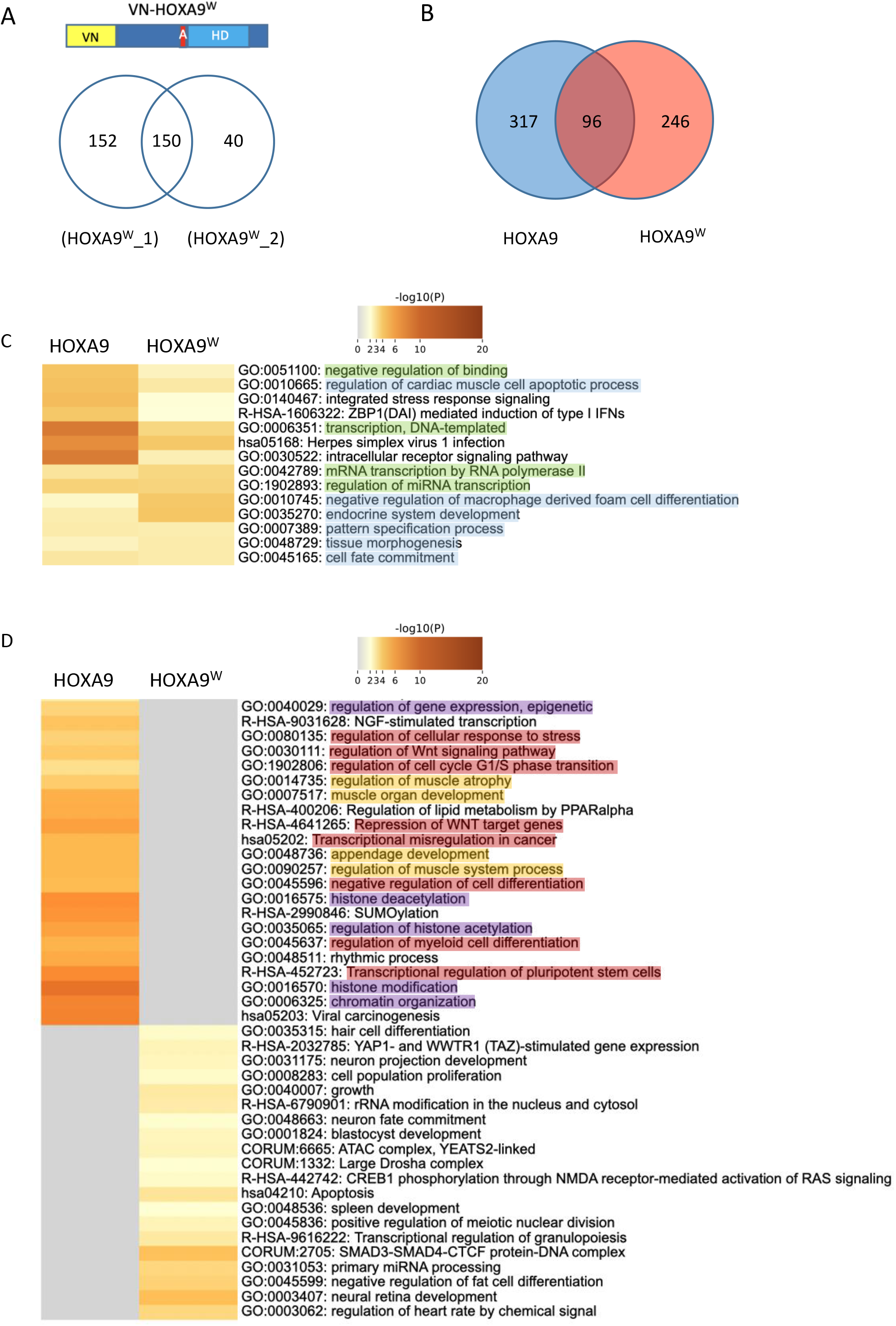
Cell-PCA reveals distinct interactomes for HOXA9 and HOXA9^W^. **A**. Venn diagram of HOXA9^W^-interacting ORFs in the two CC-HEK cell lines. The Trp (W) mutation into an Ala (A) is shown in the schematized VN-HOXA9^w^ protein above the Venn diagram. **B**. Venn diagram showing the comparison between HOXA9 and HOXA9^w^ interactomes. **C**. Heatmap for the top 20 enriched biological functions in both HOXA9 and HOXA9^W^ interactomes. One row per function, using a discrete color scale to represent statistical significance (from high (dark red) to no (gray) significance). Blue and green colors highlight functions involved in transcriptional regulation or morphogenesis, respectively. **D**. Heatmap showing the specific biological functions underlying HOXA9 and HOXA9^W^ interactomes (considering the 317 HOXA9-specific and 246 HOXA9^W^-specific interactions). Violet, orange and red colors respectively highlight functions involved in epigenetic, muscle formation and cell proliferation/cancer, which are lost upon the W mutation.

The comparison of the biological functions enriched in HOXA9 and HOXA9^w^ interactomes revealed that the classical HOX functions related to morphogenesis and DNA-binding dependent transcription were present in both interactomes, although with different levels of enrichment (highlighted in blue and green in Fig. 3C). In contrast, several functions related to epigenetic and chromatin organization were lost in the HOXA9^w^ interactome (highlighted in violet in Fig. 3D). In addition, the Trp mutation affected the activity of HOXA9 for muscle formation (highlighted in light-orange in Fig. 3D) and led to a loss of several functions involved in cell division and cancer progression (highlighted in red in Fig. 3D). This last effect has previously been reported in several studies with HOXA9 (Dickson *et al*, 2013; Ando *et al*, 2014). Finally, the ectopic functions revealed with the Trp mutation were also linked to cell division processes (Fig. 3D), suggesting that this residue could have dual activities. Other ectopic functions were related to more specific pathway such as CREB1, YAP/TAZ or ATAC complex (Fig. 3D).

Together, results obtained with VN-HOXA9 and VN-HOXA9^W^ validated the proof-of-concept ORFeome-wide interaction screen for capturing (sensitivity) and distinguishing (specificity) interactomes between two highly related HOX proteins. These results were encouraging for applying the experimental strategy in a more systematic HOX interactome screen exploration. We therefore applied the Cell-PCA strategy to capture the interactome of seven additional human HOX proteins, tackling the general issue of human HOX interactome specificity in the same cell system.

### Using Cell-PCA for a global comparison of HOX interactomes

HOX members belonging to anterior (HOXA1 and HOXA2), central (HOXC6, HOXA7 and HOXC8) and posterior (HOXA9, HOXD10 and HOXB13) paralog groups were chosen for the ORFeome-wide comparison between different HOX interactomes (Fig. 4A). HOX proteins were fused to the VN fragment, as previously described (Dard *et al*, 2018a, 2019b), and each VN-HOX encoding plasmid was transfected in the two CC-HEK-1 and CC-HEK-2 cell lines for the ORFeome-wide BiFC screen. We applied the same selection criteria as previously described with HOXA9 for sorting fluorescent cells and selecting candidate interaction partners (see Materials and methods).

**Figure 4.**
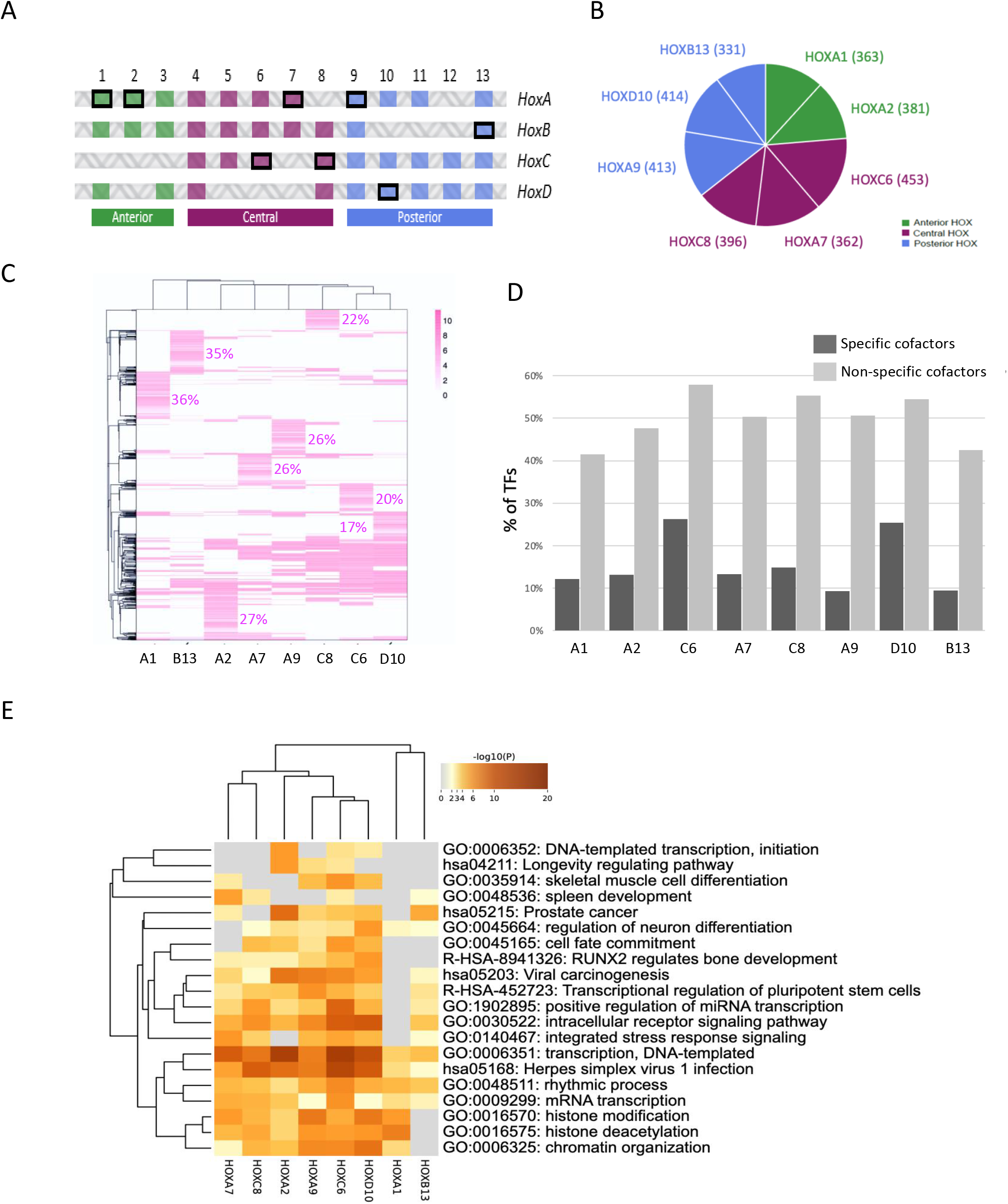
Application of Cell-PCA for global HOX interactome screening and comparison. **A**. Schematic arrangement of the 39 human HOX genes. HOX genes belong to anterior (green) central (purple) and posterior (blue) paralog groups. The 8 HOX genes used in the screen are highlighted (framed in black). **B**. Pie chart illustrating the number of interacting ORFs identified for each HOX protein in the Cell-PCA screens. **C**. Heatmap of interacting ORFs identified for each HOX protein in the Cell-PCA screens. Hierarchical clustering was performed on both column and row, according to the Pearson distance based on log2 fold change (FC) values, using average method. The proportion of HOX-specific interactions is indicated (as a % of total interactions for each HOX protein). Scale bar indicates enrichment score (log2FC) for each HOX-interacting ORFs. The proportion of HOX-specific interactions is indicated. **D**. Distribution of DNA-binding-domain containing transcription factors (TFs) among HOX-specific (dark gray) or non-HOX-specific (light gray) interactors. Note that TFs constitute between 50% and 80% of the total interactions but they are systematically more enriched in the non-HOX-specific category. **E**. Heatmap of the top-20 enriched functional profiles of the different HOX-interacting proteins. Hypergeometric p-values and enrichment factors were calculated and used for filtering. A hierarchical clustering was performed on both column and row based on Kappa-statistical similarities among their gene memberships. A discrete color scale is used to represent statistical significance. Gray color indicates a lack of significance. The large majority of these functions is linked to gene regulation or cell differentiation and development, as expected.

Results showed that each HOX members had a variable number of positive interactions (between 4.9% and 6.7% of positive interactions, Fig. 4B and Datasets EV5). The majority of these interactions are not unique, being also found with one or more additional HOX proteins (between 64% and 83%: Fig. 4C). Still, each HOX protein showed a specific cluster of interactions (between 17% and 36%: Fig. 4C). Interestingly, HOX-specific clusters are poorly enriched in DNA-binding-domain (DBD)-containing TFs (between 9% and 26%: Fig. 4D). In contrast, DBD-containing TFs are significantly more enriched in non-specific HOX fractions (between 41% and 58%: Fig. 4D).

As expected, the heatmap of the top-20 enriched functions among all HOX interactions revealed functions linked to transcriptional regulation, chromatin organization, cell differentiation and tissue/organ morphogenesis (Fig. 4E). Surprisingly, prostate cancer was also specifically enriched for several HOX proteins although it corresponds to a deregulated biological function. (Fig. 4E). Even more surprisingly, this function was most enriched for HOXA2 although prostate cancer is more often associated to the deregulated activity of posterior HOX members (see below). However, recent work showed that HOXA2 was associated with aggressive prostate cancer, underlining the robustness of our data (Gao P, et al., 2018; Xia JH, et al., 2018; Song YP., et al., 2022).

The analysis of the interactome involved in mRNA transcription confirmed that the majority of interactions established between HOX proteins and TFs were not specific (Fig. S6). Instead, it is the combination of the full set of interactions established by each Hox protein that is specific (Fig. S6). This observation suggests that HOX transcriptional specificity results principally from specific combinations of interactions rather than from interactions with specific TFs.

Altogether, results obtained showed that the Cell-PCA approach was efficient for revealing specific interactomes of eight different HOX proteins in the same cell context.

### CREB3L4 interacts with HOXB13 and promote its proliferative activity in a prostate cancer cell context

Prostate cancer was among the top-20 enriched biological functions revealed upon the analysis of HOX interactomes. The global involvement of HOX proteins in human cancer is well-established, with pro-or anti-tumoral activities, depending on the HOX protein and the cancer type (Joy Jonkers and Priya Pai, 2020; Yangyang Feng et al., 2021). More particularly, several HOX proteins have been described to promote or inhibit prostate cancer progression (Chen et al., 2012; Morgan et al., 2014). These studies rely on the analysis of the HOX expression level in primary prostate cancer cells, and functional readouts in established prostate cancer-derived cell lines. Along this line, one of the best case-study in prostate cancer is HOXB13, which has been described in several instances to be both overexpressed and required for prostate cancer cell proliferation and metastasis (Huang et al., 2014; Calvin VanOpstall et al., 2020). We therefore looked more precisely at the HOXB13 interactome involved in prostate cancer and found several interesting candidates known to display the same pro-oncogenic activity (Fig. 5A).

**Figure 5.**
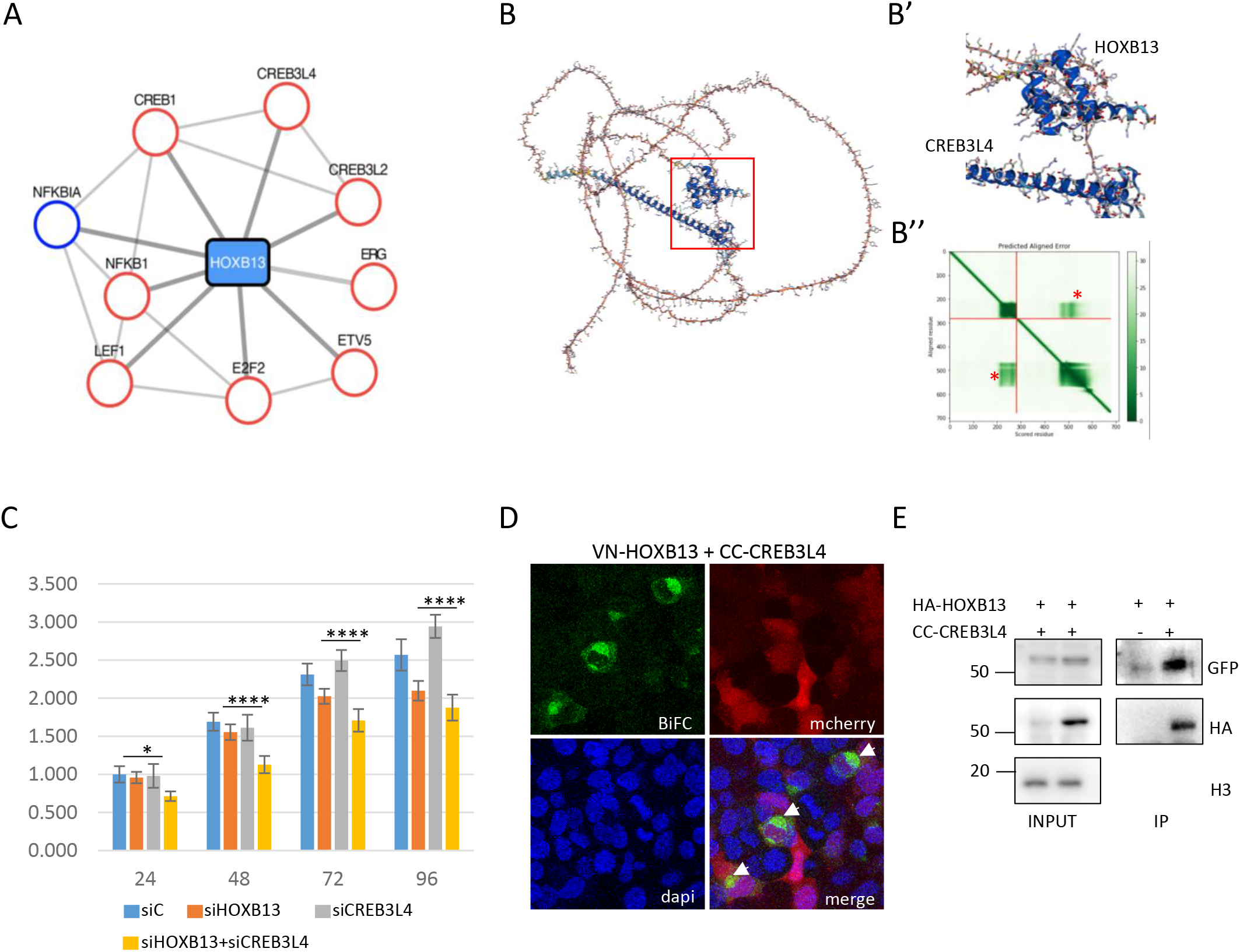
CREB3L4 is a novel cofactor of HOXB13 for promoting the proliferation of prostate-cancer derived PC-3 cells. **A**. Interactome of the HOXB13 enriched cluster involved in prostate cancer. Candidate cofactors have a known tumor suppression gene (TSG, blue circle) or oncogenic (red circle) function. **B**. AlphaFold prediction of the interaction between CREB3L4 and HOXB13. **B’**. Enlargement on the interaction interfaces between the homeodomain (HD) of HOXB13 and the alpha-helix of CREB3L4. **B”**. Prediction score of intra-domain interactions for HOXB13 (upper dark green box, corresponding to the HD), CREB3L4 (lower dark green box, corresponding to the alpha-helix) and extra-domain interactions between the HD of HOXB13 and the alpha-helix of CREB3L4 (green boxes highlighted by a red star). The first 0-280 residues correspond to HOXB13, the following 280-700 residues correspond to CREB3L4. **C**. Xcelligence assay of PC-3 cells transfected with the different siRNAs, as indicated. siC= siRNA control (see also Materials and Methods). Measures are performed at different time points post-transfection and result from three independent biological replicates. Two-way ANOVA with Tukey’s multiple comparisons, *p<0.05 and ****p<0.0001. **D**. Illustrative confocal picture of BiFC (green) between VN-HOXB13 and CC-CREB3L4 in fixed HEK293T cells. The mCherry reporter (red) stains for transfection and DAPI (blue) for nuclei. A typical enrichment is observed in dividing nuclei (white arrows). **E**. Co-IP between HA-tagged HOXB13 and CC-CREB3L4 co-expressed in HEK293T cells. The co-IP was performed with anti-HA and CREB3L4 was revealed with anti-GFP that recognizes the CC fragment (see Materials and Methods). “+” and “−” respectively denote the co-transfection or not of HA-HOXB13 with CC-CREB3L4. The protein size scale is indicated on the left side (KDa). Confocal and western blot pictures are illustrative of two independent biological replicates.

Several model cell lines derived from prostate cancers have been established. In particular, the role of HOXB13 has been well studied in the PC-3 cell line, with a role for promoting proliferation and malignancy (Y-R Kim et al., 2014; In-Je Kim et al., 2014; Aya Misawa and Yukihiro Kondo, 2021). We hypothesized that this proliferative activity could depend on the interaction with cofactors that were revealed in our screen. In such a scenario, the loss of the candidate cofactor should affect the proliferative activity of HOXB13. To test this hypothesis, we selected *CREB3L4*, which has been described to be expressed in several prostate-cancer derived cell lines, including the PC-3 cells (Uta Schmidt et al., 2006). Interestingly, CREB3L4 and HOXB13 are predicted to interact when using AlphaFold (Fig. 5B-B” and (Jumper *et al*, 2021)(Varadi *et al*, 2022)). The role of this candidate cofactor on HOXB13 proliferative activity was tested by using the xCELLigence system, which uses cellular impedance to continuously measure the number, size and surface attachment strength of adherent tumor cells (Cerignoli F. et al., 2018; Şimşek S. et al., 2020; Tribollet V. et al., 2022). We first confirmed that *HOXB13* and *CREB3L4* were expressed in PC-3 cells and that the siRNAs were efficient in affecting the corresponding endogenous gene expression (Fig. S7). Effects of the siRNAs were analyzed 24h, 48h, 72h and 96h post-transfection. The effect of each individual siRNAs against *HOXB13* or *CREB3L4* was first apparent at 72h. These effects were moderate and similar for each siRNA and more pronounced at 96h post-transfection (Fig. 5C). Importantly, combining the two siRNAs against *HOXB13* and *CREB3L4* led a significantly more pronounced effect on cellular impedance than each single siRNA, already at 24h post-transfection (Fig. 5C). Given that the siRNAs against *CREB3L4* did not affect the expression of *HOXB13* (Fig. S7), we concluded that HOXB13 and CREB3L4 could work as a cooperative dimeric complex protein for promoting the proliferation of PC-3 cells. This hypothesis was confirmed by doing BiFC and co-IP experiments that showed that HOXB13 and CREB3L4 could indeed interact and form a protein complex (Fig. 5D and E).

Altogether, these results place CREB3L4 as a new partner of HOXB13 for promoting its proliferative activity in prostate-cancer derived PC-3 cells.

## Discussion

### Cell-PCA allows capturing and comparing interactomes in the same live cell context

Protein interactomes are versatile networks involving hundreds of transient and low affinity interactions. Over the last years, several experimental strategies based on PCA systems have been developed to capture these molecular interactions, leading to promising alternatives in addition to LC-MS or Y2H-based approaches. In this context, BiFC-based PCA is particularly well-adapted for revealing pair-wise interactions in live cells and has been applied in several screening strategies to study interactomes of various bait proteins (Remy & Michnick, 2004; Berendzen *et al*., 2012; Ding *et al*, 2006; Miller *et al*, 2015). Although these screens were based on an off/on readout, with no enrichment scores neither any replicate, they revealed specific sets of interactions that were further confirmed by alternative molecular and functional assays. Altogether, this previous work established BiFC as a powerful method for revealing novel interactomes. Our work further enriches the repertoire of applicability of BiFC for large-scale protein interaction screens, in particular by proposing an experimental setup that allows using the same cell line for performing different screens. This strategy provides an additional level of information that allows comparing interactomes of different bait proteins.

Cell-PCA relies on the establishment of a cell line that has integrated a BiFC-compatible ORFeome. This cell line can be amplified and used several times for BiFC interaction screens with different bait proteins. As a proof of concept, we used the HOX protein family and two different CC-HEK cell lines, and proposed stringent filtering parameters for selecting most relevant candidate interaction partners. The screen was voluntary performed in conditions of low expression level for each CC-ORF prey construct, allowing getting specific BiFC signals with the transfected VN-bait protein (that is itself under multiple copies).

The two CC-HEK cell lines were used as biological replicates by considering the common pool of randomly integrated CC-ORFs. We also noticed that a higher number of positive interactions were systematically revealed with the CC-HEK-1 cell line when compared to the CC-HEK-2 cell line. It was probably due to better transfection conditions in the CC-HEK-1 cells (which is an inherent part of variability when considering two different biological replicates). Still, we found a relevant proportion of reproducibility of positive interactions when considering the common pool of integrated genes in the two CC-HEK cell lines, from 7% to 11% depending on the HOX protein (Fig. S8). This score is below the rate of reproducibility described for approaches based on high-throughput mass-spectrometry protein complex identification (around 19% when considering proteins present in two datasets: (von Mering C. et al., 2002) or Y2H (around 20%, (Brückner A. et al., 2009)), which can be explained by the random insertion of each CC-ORF. In particular, variation in the frequency (% of cells having integrated the CC-ORF) and expression level (depending on the genomic insertion site) between the two CC-HEK cell lines could influence the final enrichment in the fluorescent cell population. This point suggests that the level of reproducibility will probably be higher between replicates performed with the same CC-HEK cell line.

The specificity of the tools was confirmed by the results obtained with HOXA9 and the mutated HOXA9^W^ construct, with a small proportion of common interactions (23% of HOXA9 interactions were also found with HOXA9^W^). Along the same line, we got only 15 interactions that were common to all the tested HOX proteins (all are nuclear proteins with a majority of DBD-containing TFs: Fig. S9), which confirmed that each sorted BiFC-positive cell was the result of specific interactions with the HOX protein and not with the VN tag.

Nevertheless, having a neutral VN bait protein could also help in selecting most relevant interactions over the non-specific background, especially when testing few bait proteins or in the absence of a negative control. It could be the VN tag alone or fused to a fluorescent protein like mCherry.

Finally, positive interactions that were randomly pick-up from the screen with HOXA9 were subsequently validated by doing individual BiFC and Co-IP experiments. Co-IPs were performed independently of the complementation system, and were systematically reproducing the BiFC result. Similar observations have been reached from previous BiFC screens in cells (Lee *et al*, 2011)(Sung *et al*, 2013) or in the *Drosophila* embryo (Baëza *et al*, 2015), underlining that BiFC and Co-IP can be used as complementary approaches for validating an interaction potential between two artificially expressed candidate proteins. Our results also validated our stringent selection criteria as appropriate for selecting relevant candidate interactions.

The Cell-PCA approach did not reveal all expected or previously described interactions. For example, we found that not all HOX proteins were able to interact with the representatives of the generic PBX and MEIS cofactors present in the two CC-HEK cell lines (PBX3 and MEIS2). PBX3 was captured for HOXC6 and HOXA7, and MEIS2 for HOXA1 and HOXC8. Our stringent selection criteria eliminated HOXA1, HOXA2, HOXA9, HOXC10 and HOXB13 for PBX3, and HOXC6, HOXA9, HOXD10 and HOXB13 for MEIS2 (0 read was obtained in the two CC-HEK cell lines for HOXA2 and HOXA7 with MEIS2). These results suggest that the fusion topology might not be best appropriate in several cases for revealing the interactions by BiFC. Performing a second screen with alternative fusion topologies could help in resolving this issue. Alternatively, the *CC-PBX3* and *CC-MEIS2* constructs could also be be weakly expressed in most cells of the CC-HEK cell library for different reasons (insertion sites, post-translational modifications, etc.). Along the same line, the random insertion of each CC-ORF construct can lead to a bias that we do not control for positive binding partners: the frequency of interaction with the HOX protein will also depend on each cell-specific level of expression of the CC-ORF within the cell population. Using CRISPR-based system for targeting a unique genomic insertion site for all CC-ORF constructs could constitute an interesting alternative with this regard in the future (Chi, X. et al., 2019).

Although further developments could be done in the future, our results show that Cell-PCA is a sensitive, specific and robust approach for performing ORFeome-wide interaction screens upon simple transfection of a bait protein. Cell-PCA not only simplifies the protocol (the screen can be performed in a classical A2 laboratory environment since it does not rely on systematic transduction, as previously described (Ding *et al*, 2006)), but also enables testing different bait proteins in the same batch of cells, therefore providing a unique level of information for comparative interactome analyses.

### HOX interactomes revealed by Cell-PCA: molecular properties and comparison with existing databases

The analysis of eight different HOX interactomes revealed several unexpected and interesting molecular features. For example, there was an important proportion of non-TFs in the overall pool of HOX interacting proteins. Similar observations have been raised from a Y2H screen with HOXA1 (Lambert *et al*, 2012; Taminiau *et al*, 2016). Along the same line, the Hox protein Ultrabithorax (Ubx) has been shown to establish tissue-specific interactions with cofactors involved in translational regulation (Carnesecchi *et al*, 2020). Altogether, this novel layer of interactions illustrates the ability of HOX proteins to be engaged in the regulation of several post-transcriptional regulatory processes, a level that remains poorly investigated to date.

Interestingly, TFs were mostly found in non-HOX specific interactions, whereas the proportion of non-TFs was enriched in HOX-specific interactions. Accordingly, HOX interactomes related to transcriptional regulatory processes contained distinct combinations of a majority of non-specific interacting TFs (Fig. S6). This observation underlines that HOX transcriptional specificity mostly relies on the establishment of specific combinations of interactions with TFs that have the potential to interact with several HOX proteins. This molecular mode of action has already been proposed for *Drosophila* Hox interactomes, suggesting that it could be a general and conserved feature underlying Hox transcriptional specificity (Baëza *et al*, 2015).

We also compared our positive interactions to the publicity available Biogrid database (https://thebiogrid.org/), which compiles all characterized protein-protein interactions with genetic and/or molecular evidence in different model systems. In particular, Biogrid includes interactions listed in Bioplex (https://bioplex.hms.harvard.edu/). Bioplex corresponds to interactions obtained with the ORFeome v8.1 expressed as HA-FLAG-tagged bait proteins for doing co-IP in different human cell lines, including HEK293T cells, followed by LC-MS identification of endogenous binding partners.

In general, we found between 5% and 7% of the 6713 integrated genes as positive HOX interaction candidates in our BiFC screens. In comparison, a Y2H screen performed with HOXA1 against the human ORF v3.1 revealed only 59 positive interactions, therefore less than 1% of the screened bait proteins (Lambert *et al*, 2012). This result illustrates one major drawback of the Y2H heterologous system, with a lack of sensitivity and a number of false negatives generally obtained in Y2H screens (Brückner A. et al., 2009). Nevertheless, considering all positive HOXA1 interactions listed in Biogrid from different Y2H screens revealed a total number of 347 positive interactions, therefore around 3.5% of the total human ORFeome. Cell-PCA revealed 5.4% positive interactions for HOXA1 (363/6713), which is in the same order of magnitude. The higher proportion of positive interactions found with Cell-PCA could be explained by the more appropriate (with natural DNA-binding of HOXA1 in a human live cell context) therefore more sensitive biological system than Y2H. Still, we found that 71 proteins out of the 254 captured interactions in Y2H screens with HOXA1 and present in our CC-HEK cell lines were also positive with HOXA1 in our BiFC screen, making a highly relevant percentage of overlap (28%, Fig. S10). In contrast, only 5/347 Y2H HOXA1 positive interactions are also listed in Bioplex. It is interesting to note that 2/5 of these listed Bioplex cofactors were present in our CC-HEK cell lines and that one of it has been captured with VN-HOXA1 (ZNF503). In any case it is important to stress out that, in contrast to cell-type specific co-IP of endogenous cofactors, Y2H-or BiFC-based approaches are revealing a global potential of interaction between the bait and the candidate interaction partner.

The comparison with interactions listed in Biogrid for HOXC8 showed the same range of overlap: among the 119 listed cofactors for HOXC8, 98 were present in our CC-HEK cell lines and 29 were positive (29.6%, Fig. S10). Much less cofactors were described for the other HOX proteins in Biogrid, which limits the scope of the conclusions. For example, 20/27 listed cofactors of HOXA9 were present in our CC-HEK cell lines, and 7/20 were positive in our BiFC screens, making a similar range of positive percentage (35%, Fig. S10). Overall, these observations underline that interactions revealed with Cell-PCA were found at a significant proportion in the current interactomics database (which result mostly from Y2H screens).

In conclusion, our work confirmed that HOX proteins have a strong potential to engage a number of interactions with various partners, and in particular partners involved in so far poorly investigated post-transcriptional regulatory processes. Our understanding of their specific molecular mode of action will certainly require a better consideration of these supplementary levels of regulation in appropriate cells or tissue-systems in the future.

## Supplementary Figures

**Supplementary Figure 1. Application of PCA-based strategies for large-scale interaction screens in living cells**. The co-transfection PCA-based screening strategy (Co-PCA) relies on the transitory expression of the FPN-fusion bait protein and the FPC-fusion human ORFeome library upon co-transfection in the cell line (Remy & Michnick, 2004; Berendzen, K.W. et al., 2012) (B). The retrovirus PCA-based screening strategy (Re-PCA) relies on the use of a cell line stably expressing the FPN-fusion bait protein and infected by retroviruses containing the FPC-prey human ORFeome library that is artificially expressed with a constitutive promoter (Lee *et al*, 2011; Simon E. Coopera, 2015) (D). A first generation of the Re-PCA screening strategy was based on random genomic insertion of retroviruses encoding the FPC in the three possible open reading (Ding *et al*, 2006).

**Supplementary Figure 2. Map of the lentiviral expression vector and basal expression level of the CC-ORF constructs in the established CC-HEK cell line. A**. A lentiviral destination vector (pLV-CC-Gateway) was generated to clone the human ORFeome library in frame with the C-terminal fragment of Cerulean (CC), leading to the pLV-CC-ORF vector. The expression of the CC-ORF is under the control of a first-generation TRE promoter. This system (Tet-Off) requires the co-expression of a Tet transactivator (tTA) for full activation of the promoter. **B**. The basal activity of the TRE promoter (without adding Dox and the tTA-coding vector) is sufficient to detect the expression of the inserted CC-ORFs in the CC-HEK1 cell population. Control with the original HEK-293T cell lines leads no staining. Immunostaining was performed with an anti-GFP recognizing the CC fragment (green). DAPI (blue) stains for nuclei. Scale bar = 50 μm.

**Supplementary Figure 3. Schematic of the ORF Capture-Sequencing method**.

Libraries were constructed using our own designed proprietary protocol in order to enrich in sequences covering the beginning of all the CC-ORFs inserted in the genome. Roughly, the DNA extracted from sorted cells (1) is fragmented using the Covaris S220 Ultrasonicator (2). Then, the Ion adapter P1 is non-directionally ligated to both ends of the DNA fragments according to the standard blunt-end ligation protocol for Ion Torrent libraries (3). Because the P1 libraries molecules containing hORF sequences are underrepresented, an enrichment by a first PCR is performed by using a biotinylated forward primer located on the plasmid sequence upstream the hORFs and a reverse primer located at the end of the P1 adapter (4). Biotinylated PCR fragments, containing now at each end the beginning of the hORFs and the P1 adapter respectively, are then captured using streptavidin-coupled magnetic beads (5, 6). These molecules are used as a template for the final hemi-nested PCR amplification using a forward fusion primer containing a barcoded Ion A adapter and the same reverse primer as before (7). The hemi-nested PCR reinforces the specificity of the fragments to be sequenced since the primer fused to A and the barcode is located closer to the hORF beginning on the plasmid. After a size selection using SPRI beads to meet Ion Torrent requirements (8), the qualified and quantified barcoded libraries are multiplexed in an equimolar manner and sequenced on the Ion Proton sequencer using a P1 chip following the manufacturer’s recommendations (9).

**Supplementary Figure 4. Testing for specific BiFC signal and FACS gate for BiFC-positive cell population. A**. Schematic representation of the cold (non-transfected) CC-HEK cell line. **A’**. Illustrative confocal picture of non-transfected cells upon excitation wave length for BiFC. Transmission light capture shows the cells. **A”**. FACs gates for the GFP channel. No fluorescent signal can be observed in the cold/non-transfected CC-HEK cell population. **B**. Schematic representation of VN-HOX9 transfection in the CC-HEK cell line. **B’**. Illustrative confocal picture of transfected cells upon excitation wave length for BiFC. Fluorescent cells are indicated (white arrows). Transmission light capture shows the cells. **B”**. FACs gates for the GFP channel. Fluorescent signals can be observed in the CC-HEK cell lines upon transfection of VN-HOXA9. Compared to A”, the specific GFP-positive population can be isolated from the negative/non-fluorescent cells. This population represents 2.67% of the total cell population. A similar percentage of fluorescent cells (between 1,5% and 3%) was obtained with the different HOX constructs.

**Supplementary Figure 5. Individual validation of positive HOXA9 interactions selected from the BiFC screen in the two CC-HEK cell lines**. Illustrative confocal pictures of BiFC and co-immunoprecipitation (Co-IP) in HEK293T cells between HOXA9 and the selected candidates, as indicated. Confocal and western blot pictures are illustrative of at least two independent biological replicates. For BiFC experiments, the mCherry reporter (red, merge panels) is indicative of the transfection efficiency. Note the various intra-cellular BiFC profiles with different candidates. Scale bar, 10μm. Co-IP was performed with anti-HA recognizing HA-HOXA9 and the CC-ORF was revealed with anti-GFP. Staining with Histone H3 antibody validates the correct protein extraction in each condition. “+” and “−” respectively denote the co-transfection or not of HA-HOXA9 with the CC-ORF construct. The protein size scale is indicated on the left side (KDa).

**Supplementary Figure 6. Interactome of the enriched RNA-PolII dependent transcription function found in HOX BiFC screens**. Transcription factors are boxed in a rectangle whereas non transcription factor partners are boxed in a rhomb. Yellow-colored boxes represent specific interactors of one HOX protein. Anterior HOX are in green, central HOX are in purple and posterior HOX are in blue.

**Supplementary Figure 7. Expression of *HOXB13, CREB3L4* and *ERRα* 48h post-transfection of the indicated siRNAs in PC-3 cells**. Each siRNA significantly reduces the expression of its corresponding gene as evaluated by rt-qPCR. Results were obtained from eight independent experiments. Bars represent mean +/-sem. **: *p*<0.01, ***: *p*<0.005. t-test values (relative to control) are shown for significantly different points.

**Supplementary Figure 8. Reproducibility of positive interactions among the 5005 commonly integrated CC-ORFs in the CC-HEK-1 and CC-HEK-2 cell lines**. The third column is indicative of the percentage of reproducibility (% of commonly integrated CC-ORFs that were positive in both CC-HEK1 and CC-HEK2 cell line screens).

**Supplementary Figure 9. List of interactors that are positive with all tested HOX proteins**. All these interactors are nuclear proteins. DNA-binding-domain containing TFs are highlighted (bolded).

**Supplementary Figure 10. Comparison of Hox positive interactions between Cell-PCA and the Biogrid database**. Only genes corresponding to integrated CC-ORFs in the CC-HEK cell lines have been considered for the comparison. Paralogs and isoforms have also been considered to increase the number of genes that could be analyzed.

## Supporting Information

**Dataset EV1. List of 5799 ORFs expressed in CC-ORFeome HEK cell line-1**.

**Dataset EV2. List of 5549 ORFs expressed in CC-ORFeome HEK cell line-2**.

**Dataset EV3. List of HOXA9-interacting proteins identified by Cell-PCA screening**.

**Dataset EV4. List of HOXA9**^**W**^**-interacting proteins identified by Cell-PCA screening**.

**Dataset EV5. Full list of HOX-interacting proteins identified by Cell-PCA screens**.

**Dataset EV6. hORFs identified with the first 50bp (with up to 3 bp differences) upon**

**BLASTN with the hORFeome v3.1**.

## Materials and Methods

### Cell Lines

HEK-293T and PC-3 cells were purchased from Europen Collection of Authenticated Cell Cuture (ECACC) through the biological resource center Anira-AGC platform of the SFR Biosciences UAR3444/US8 of Lyon. Both cell lines were cultured in Dulbecco’s modified Eagle’s medium (DMEM-GlutaMAX-I, Gibco by Life Technologies) supplemented with 10% (v/v) heat inactivated fetal bovine serum (FBS) and 1% (v/v) Penicillin-Streptomycin (5,000U penicillin and 5mg streptomycin/mL), incubating at 37°C, in an atmosphere of 5 % CO_2_. HEK-293T-CC-ORFs were cultured as above with 0.3 µg/ml of puromycin (Gibco, Cat No. A1113803) in their culture medium.

### Plasmids

The bait plasmids pcDNA3-VN-HOXs (expressing VN-HOXs) were made as described previously (Dard *et al*, 2019b, 2018b). The lentiviral pLV-CC-ORFs vector collection was kindly provided by P. Mangeot (CIRI, ENSL, France). The genomic DNA was used for library preparation and subjected to next-generation sequencing at in-house NGS sequencing platform (PSI, IGFL, Lyon, France). About 8 200 ORFs from the V3.1 version of the hORFeome were fused at the 5’ end to the C-terminal part of the mCerulean gene (encoding the last 155-238 aa) using the Gateway® technology.

For individual BiFC tests, constructs were cloned into the pLIX _403 vector (a gift from David Root, Addgene plasmid # 41395; http://n2t.net/addgene:41395; RRID:Addgene_41395). DNA sequencing of all constructions were carried out at GENEWIZ Company (Germany). All vectors are available upon request.

### Lentivirus Preparation and Infection

The pooled lentiviral constructs pLV-CC-ORFs were packaged into lentivirus particles at the AniRA-Vectorology core facility (SFR Biosciences UAR3444/US8, Lyon, France). HEK-293T cells were transduced in independent replicates with two batches of lentivirus (CC-ORF library 1 and CC-ORF library 2) at a low multiplicity of infection (0.3) to achieve approximately one-gene-one-cell condition(Yea, K., Zhang, H., Xie, J., Jones, T.M., Yang, G., Song, B.D., and Lerner, 2013), with ≥500X representation. Culture medium was supplemented with 8 µg/mL polybrene (Sigma) at the time of transduction, and was changed the next day. Two days after transduction, cells were selected with 0.5 µg/mL Puromycin (Gibco, Cat No. A1113803) for 4 days, until the uninfected control cells completely died and the selected cells reach near confluency. Final amplified transduced cells were split into aliquots of 4×10^6^ cells each and stored in liquid nitrogen for future screens.

### Immunostaining of CC-ORF transduced HEK cells

1×10^5^ cells were seeded on glass coverslips in 24-well plates. Twenty-four hours after plating, cells were fixed in 4% paraformaldehyde in PBS for 15 min at room temperature, permeabilized in 0.3% Triton X-100 for 10 min, and rinsed in PBS. The cells were preincubated in 3% Bovine serum albumin-PBS at room temperature for 1 h and incubated overnight at 4°C with primary antibodies against CC-ORF (rabbit anti-GFP, polyclonal, Invitrogen A11122, 1:1000). The sections were then incubated with the corresponding fluorescein-conjugated secondary antibodies (Alexa Fluor 488 anti-rabbit, Invitrogen A11008, 1:1000) for 2 h at room temperature. Coverslips were mounted in VECTASHIELD Antifade Mounting Medium with DAPI (VECTOR, Cat No. LS-J1033-10). They were analyzed via confocal microscopy (Zeiss LSM780).

### Cell-PCA Screen

8.10^6^ CC-HEK cells (~800X representation) were thawed and passaged for 2 population doublings to recover. For each screen, aliquots of 6 ×10^6^ recovered cells were seeded in two 6-well plate (500k cells/well) and grown for 24 hours for achieving a final confluence around 80%. The basal expression of the TRE promoter was used for the expression of the CC-ORFs in the cell population. The transfection of different bait plasmids pcDNA3-VN-HOXs was performed using the jetPRIME reagent (Polyplus, Ref 114-15) following manufacturer’s instruction. After 18 h of transfection, all transfected cells were pooled and BiFC-positive cells were subsequently sorted using a BD FACS Aria II Cell Sorter (AniRA-Cytometry core facility of the SFR Biosciences UAR3444/US8, Lyon, France). After each screen, sorted cells were harvested and genomic DNA was extracted using PureLink Genomic DNA mini kit (Invitrogen, Cat No. K182001), according to manufacturer’s instruction. The genomic DNA was used for library preparation and subjected to next-generation sequencing at in-house NGS sequencing platform (PSI, IGFL, Lyon, France).

### Next Generation Sequencing and identification of the positive hORFs

Libraries were constructed using our own designed proprietary protocol in order to enrich in sequences covering the beginning of all the hORFs inserted in the genome (see details in Fig. S2). After a size selection using SPRI beads to meet Ion Torrent requirements, the qualified and quantified barcoded libraries were multiplexed in an equimolar manner and sequenced on the Ion Proton sequencer using a P1 chip following the manufacturer’s recommendations.

NGS raw data were analyzed with the Galaxy instance (Afgan, E., et al. 2018) of the ENS of Lyon and maintained by the Centre Blaise Pascal (CBP, ENS Lyon). A dedicated Galaxy pipeline was created to identify the hORFs detected by sequencing and their associated read counts for each barcoded sample. After demultiplexing, the raw reads were trimmed to remove very low-quality bases in the 3’ and 5’ ends using a sliding window process. By construction, the libraries are oriented and all the reads begin with the same short plasmid sequence that is present upstream of any inserted hORF. This sequence was removed by Cutadapt (Martin, 2011), by allowing a maximum error rate of 0.15. Only the trimmed reads beginning with ATG were retained for further analysis. This last step removes any reads that could result from a non-specific PCR amplification. Because read length is variable with the Ion Torrent technology, the reads were then trimmed to 50bp to have all the same length. These reads were next compared to the hORFeome.V3.1 database by BLAST tool (Cock P.J.A., et al. 2015) using strict conditions (only one hit retained with at least 98% of identity on 95% of the query coverage; matches starting to position 1 of the hORFs). For each hORF that obtains hits, the number of reads matching this hORF was counted and sum up in a table for further analyses. According to the criteria used, only hORFs that have more than 3 bases of difference in their first 50bp can be differentiated (98% of the genes of the hORFeome can still be distinguished, dataset EV6). Moreover, when the beginning of the hORFs is identical or nearly identical (mainly hORFs corresponding to different isoforms of the same gene), the read is assigned to only one of the possible alternatives, usually always the same. Since the analyses were done at the gene level, the counts of ORFs for a same gene were add-up.

### Identification of candidate HOX-interacting ORFs

After raw data cleaning, the sequencing counts for each gene in each library were normalized to 10M. To limit artefacts, count tables were denoised using an arbitrary threshold of at least 50 counts/10M by gene in the cold cell-control libraries. For the sorted cells, a higher threshold of 500 counts (x10) was applied considering they constitute an enriched population when compared to non-sorted/cold cells. Then, the number of counts in sorted cells was divided by the number of counts in the cell control libraries and the log2FC was calculated as an enrichment score (ES) for each gene.

To generate the final list of HOX-interacting candidates, genes that were present in both replicates with an ES>= 1.5 were combined, assigning the highest ES obtained to each gene. To get an extensive view of all potential interactions, we also considered the top-enriched genes (with a stringent ES>=6) for interactions present in a single replicate.

### Individual BiFC validation in live cells

For transfection, 3.10^5^ cells were seeded on glass coverslips in 6-well plates and incubated for 24h. Then, cells were transfected with jetPRIME (Polyplus, Ref 114-15) following manufacturer’s instruction. A total of 1.75ug of plasmid DNA were transfected per well: 750 ng of plix-VN-HOXA9, 750 ng of plix-CC-ORF and 250 ng plix-mCherry plasmids. After 18 hours of incubation in the presence of doxycycline (100 ng/ml final), the cell-coated coverslip was taken and mounted carefully on a glass slide for image capture under confocal microscopy (Zeiss LSM780). All samples were imaged using identical settings and quantified as previously described (Dard *et al*, 2018b). Two biological replicates were systematically done, using the interaction between HOXA9 and PBX1 as a positive BiFC control and the mCherry reporter for assessing transfection efficiency.

### Individual Co-immunoprecipitation (IP) validation

For co-IP assays, HEK-293T cells were plated at 2 million cells in a 10cm-petri dishe and transfected with either pLIX_403-2HA-HOXA9 (4µg) and/or pLIX_403-CC-bait (4µg) with PEI at a ratio N/P=5. Cells were returned in a complete medium supplemented with 200ng/ml of doxycycline to induce expression of the bait and the preys. Cells were harvested 48h-post transfection in Phosphate Buffered Saline (PBS) and pellets were resuspended in NP40 buffer (20 mM Tris pH 7.5, 150 mM NaCl, 2 mM EDTA, 1% NP40) and treated with Benzonase (Sigma). Anti-HA magnetic beads (Pierce) were added to the protein extract, incubated for 2 hours and washed five times with NP40 buffer. All samples were resuspended in Laemmli buffer for immunoblotting analysis. All buffers were supplemented with protease inhibitor cocktail (Sigma), 1 mM of DTT and 0.1 mM PMSF. Input fractions represent 1–10% of the immunoprecipitated fraction.

For western blot analysis, proteins were resolved on 12% SDS-PAGE, blotted onto PVDF membrane (Millipore) and probed with specific antibodies after saturation. The antibodies (and their dilution) used in this study were Histone 3 (1791 Abcam, 1/10,000), GFP (A11122 Life Technologies, 1/2000), HA (901513, Biolegend, 1/3000e) and HOXB13 (PA5-98698 Life Technologies, 1/1000).

Developing was performed using chemiluminescence reaction (ECL, GE-Healthcare) with secondary coupled to HRP (Promega, 1/5000e) and the Amersham ImageQuant 800 (Cytiva).

### Functional enrichment and Interactome analysis

Both functional and interactome analyses were performed with Metascape (https://metascape.org/) (Zhou *et al*., 2019) using custom analysis settings. Subsequently, Cytoscape v3.8.2 (Shannon *et al*, 2003) was conducted to visualize representative HOX functional interactomes.

In functional enrichment analysis, the HOX-interacting protein candidates were searched against GeneOntology Biological Processes, KEGG pathways, CORUM and Reactome databases. A p-value cutoff ≤0.01 was used to determine significant functional terms. They were then hierarchically clustered into a tree based on Kappa-statistical similarities among their gene memberships. A kappa score ≥0.3 was applied as the threshold to cast the tree into term clusters. We selected the term with the best p-value within each cluster as its representative term and display them in the heatmaps.

Physical PPIs from multiple data sources were captured for construction of the interaction networks. Homo sapiens were selected as the organism for subsequent analysis. Min network size 3 was regarded as cut-off criterion for network visualization and disconnected nodes was hidden. The complex identification algorithm, MCODE (Bader, G.D., and Hogue, 2003), was used to identify highly interconnected clusters in the network. The most important protein complex clusters in the PPI network were extracted, with default settings in MCODE, degree cutoff = 2, node score cutoff = 0.2, Max depth = 100, and k-score = 2. For each complex, it further applied function enrichment analysis and used significantly enriched terms for annotation of its biological roles. Following manual curation, the similar terms were combined and classified into non-redundant parent functions and categories, which was visualized by heatmap. Based on combined data set, all representative HOX functional interactomes were generated and visualized by Cytoscape.

### Xcelligence assays

The xCELLigence system (ACEA Biosciences), which records cellular events in real time by measuring electrical impedance across microelectrodes integrated on the bottom of culture plates (E-plates), was utilized for proliferation experiments.

First, cell culture media was added to each well of 16 well E-Plates (ACEA Biosciences) for measuring background impedance. Then, 1.5 × 10^5^ of PC3 cells were transfected with a mixture of control, ERRα, HOXB13, or CREB3L4 siRNAs at 8 pmol/ml using INTERFERin (Polyplus Transfection) according to the manufacturer’s protocol. 2,5.10^4^ of transfected cells were directly seeded on E-Plate and the impedance was measured every 15 min for 96h. The impedance signal is proportional to the number of cells proliferating in each well and displayed as Cell Index. Data are analyzed with the RTCA Software 2.0 and presented as mean +/ − SEM of three experiments performed in triplicate or quadruplicate.

siRNAs used in this study were from Eurogentec:

HOXB13: GUUCAUCACCAAGGACAAG and CUUGUCCUUGGUGAUGAAC

CREB3L4: CCAGUUCUCCUAUGCUCUA and UAGAGCAUAGGAGAACUGG

ERRα: GGCAGAAACCUAUCUCAGGUU and CCUGAGAUAGGUUUCUGCCUC

### siRNAs and RT-QPCR

For siRNA transfection, 3.10^−5^ cells per ml were seeded in 6-well plate and 25 pmol/ml of total siRNA were transfected with INTERFERin (Polyplus Transfection) according to the manufacturer’s recommendations. Total RNAs were extracted by the guanidinium thiocyanate/phenol/chloroform method. 1 µg of RNA was converted to first strand cDNA using the RevertAid kit (ThermoScientific). Real time qPCRs were performed in 96 well plates using the IQ SYBR Green Supermix (BioRad). Data were quantified by ΔΔ-Ct method and normalized to 36b4 expression.

Sequences of the primers used in this study:

36b4: GTCACTGTGCCAGCCCAGAA and TCAATGGTGCCCCTGGAGAT

HOXB13: CAGATGTGTTGCCAGGGAGA and TGCTGTACGGAATGCGTTTC

CREB3L4: AGCTGCCCTTTGATGCTCAT and CGGTCAGGAACAGGGTTTGA

ERRα: CAAGCGCCTCTGCCTGGTCT and ACTCGATGCTCCCCTGGATG

## Acknowledgements

We thank Dr. P. Mangeot (CIRI, Lyon, France) for providing the pLV-CC-ORF plasmid library, the AniRA-Cytometry, AniRA-Vectorology and AniRA-Genetic and Cellular Analysis platforms of the UAR3444/US8 (Lyon, France), and Emmanuel Quemener from the Blaise Pascal Center (ENSL) for computer resources (Galaxy platform). We thank Frédéric Marmigère for careful reading of the manuscript and helpful comments.

This work was supported by the CNRS, ENS-Lyon, Fondation pour la Recherche Médicale (FRM #160896), Association pour la Recherche contre le Cancer (ARC #PJA20191209567), Ligue Régionale contre le Cancer (#119030) and Centre Franco-Indian pour la Promotion de la recherche Avancée (Cefipra #5503-P). Y.J. received a PhD grant from the China Scholarship Council (grant #201708070003).

